# Utilizing an acute hyperthermia-induced seizure test and pharmacokinetic studies to establish optimal dosing regimens in a mouse model of Dravet syndrome

**DOI:** 10.1101/2023.10.03.560653

**Authors:** Jeffrey A. Mensah, Kristina Johnson, Tia Freeman, Christopher A. Reilly, Joseph E. Rower, Cameron S. Metcalf, Karen S. Wilcox

## Abstract

**Objective:** The current standard of care for Dravet Syndrome includes polytherapy after inadequate seizure control with one or more monotherapy approaches. Treatment guidelines are often based on expert opinions, and finding an optimal balance between seizure control and adverse drug effects can be challenging. This study utilizes efficacy and pharmacokinetic analysis of a second-line treatment regimen that combines clobazam and sodium valproate with an add-on drug as a proof-of-principle approach to establish an effective therapeutic regimen in a preclinical mouse model of Dravet Syndrome.

**Method:** We evaluated the efficacy of add-on therapies stiripentol, cannabidiol, lorcaserin, or fenfluramine to clobazam and sodium valproate against hyperthermia-induced seizures in *Scn1a*^*A1783V/WT*^ mice. Clobazam, N-desmethyl clobazam (an active metabolite of clobazam), sodium valproate, stiripentol, and cannabidiol concentrations were quantified in plasma and brain using liquid chromatography-tandem mass spectrometry for the combinations deemed effective against hyperthermia-induced seizures. The concentration data were used to calculate pharmacokinetic parameters via non-compartmental analysis in Phoenix WinNonLin.

**Results:** Higher doses of stiripentol or cannabidiol, in combination with clobazam and sodium valproate, were effective against hyperthermia-induced seizures in *Scn1a*^*A1783V/WT*^ mice. In *Scn1a*^*WT/WT*^ mice, brain clobazam and N-desmethyl clobazam concentrations were higher in the triple-drug combinations than in the clobazam monotherapy. Stiripentol and cannabidiol brain concentrations were greater in the triple-drug therapy than when given alone.

**Interpretation:** A polypharmacy strategy may be a practical preclinical approach to identifying efficacious compounds for Dravet Syndrome. The drug-drug interactions between compounds used in this study may explain the potentiated efficacy of some combination therapies.

**Graphical Abstract:** 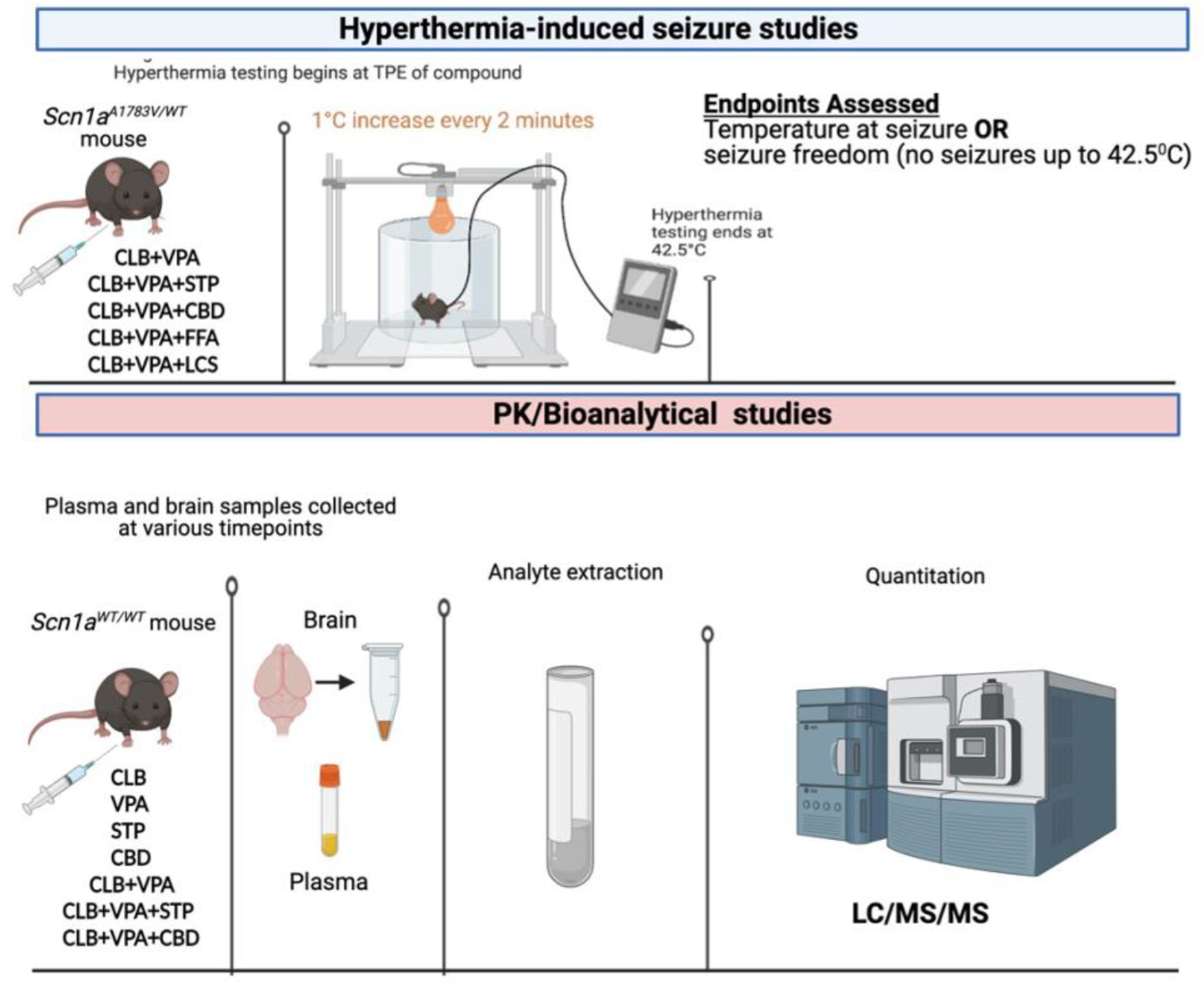

**Key findings:** The hyperthermia-induced seizure assay in *Scn1a*^*A1783V/WT*^ mice can identify efficacious antiseizure medications. A triple-drug administration paradigm was designed that mimics clinical standards for add-on therapies in DS. Stiripentol and cannabidiol were effective against hyperthermia-induced seizures when co-administered with clobazam and sodium valproate. Potential drug-drug interactions between compounds may explain the potentiated antiseizure effect in triple-drug therapies. The triple-drug “add-on” therapy approach may be a valuable preclinical strategy for screening investigational compounds for DS, an assay offered at the ETSP.

## Introduction

Dravet syndrome (DS) is a rare but catastrophic infant-onset genetic epilepsy resulting from *de novo* pathogenic *Scn1a* mutations in approximately 80% of reported cases^1,2^ Over 900 distinct mutations in the *Scn1a* gene, which encodes for the Na_v_ 1.1 ion channel, have been identified^3,4^ DS develops in the first year of life in approximately 1 per 30,000 live births.^5^ Seizure onset is typically induced by fever, which worsens and manifests as spontaneous recurrent prolonged generalized tonic-clonic seizures, sometimes resulting in status epilepticus. DS also includes developmental delays, cognitive impairment, and an increased risk for sudden unexpected death in epilepsy.

Antiseizure medications (ASMs) recommended as first- and second-line treatments and treatment regimens for DS patients have predominantly relied on expert opinions and an insufficient number of retrospective, limited sample-sized, open-label studies.^6^ Clinical response to conventional ASMs has been disappointing, with many DS patients remaining pharmacoresistant to available treatment options^.7,8^ As outlined, the treatment algorithm includes sodium valproate (VPA) and clobazam (CLB) as first-line options. Usually, as practiced in the clinic, if one first-line drug fails to offer optimal seizure control, the other agent is added on. Most pediatric DS patients use a second-line drug concomitantly with VPA and CLB.^9^ Several epileptologists have suggested stiripentol (STP) and cannabidiol (CBD) as the best second-line agents to be used in combination with VPA and CLB, with fenfluramine (FFA) emerging as a promising option^10-12^

Despite the advances in the search for effective therapies for DS, including recommended combinatorial strategies, it is difficult to find an equilibrium between seizure control, adverse drug effects, and the burden of polypharmacy in patients with DS^13,14^ The choice of the drug combination and selected tolerable doses frequently rely on the suggested treatment regimen and experience in the clinical setting.^15^ Establishing an optimal dosing regimen for DS requires understanding the relationships between dose, exposure, and effect for commonly used ASMs. This understanding hinges upon appropriate pharmacokinetic (PK) and pharmacodynamic (PD) profiling of ASMs used either as a monotherapy or combination therapy, especially during chronic treatment of DS.^16^ Preclinical studies are valuable for comparing PK with efficacy and tolerability and can inform clinical approaches. Thus, integrating PK information in an appropriate preclinical model can improve the development of polypharmacy strategies in the clinical setting.^1^

We have previously described a mouse model of DS that offers advantages for screening novel compounds using a polypharmacy approach.^17^ The present study uses this previously described mouse model to determine PK parameters and test the efficacy of clinically relevant second-line drug combination therapies. These data will enhance our understanding of PK and PD drug-drug interactions (DDI) of add-on combination therapies in the treatment of DS and generate essential data for optimizing the sub-chronic or chronic dosing regimens in a mouse model of DS.

## Materials and methods

### Animals

The Institutional Animal Care and Use Committee of the University of Utah approved all animal care and experimental procedures. Animal experiments were conducted per Animal Research: Reporting of In Vivo Experiments guidelines (https://www.nc3rs.org.uk/arrive-guidelines). We generated experimental animals by breeding a floxed stop male Scn1aA1783V (B6(Cg)-Scn1atm1.1Dsf/J, Jax #026133) with a Sox2-cre (B6. Cg-Edil3 ^Tg(Sox2−cre)1Amc^/J) female mouse to produce both heterozygous (*Scn1a*^*A1783V/WT*^) and wild-type offspring. Both female and male heterozygous and age-matched wild-type littermates were used for experiments. Mice were group-housed in a specific pathogen-free mouse facility under standard laboratory conditions (14-h light/10-h dark) and had access to food and water ad libitum, except during hyperthermia-induced seizure experiments and were transferred to the experimental room approximately 1 hour before testing. A detailed list of common data elements was recorded, and detailed case report forms were utilized to confirm all aspects of each study (available upon request).

### Hyperthermia-induced seizure test

Hyperthermia-induced seizure experiments were conducted in *Scn1a*^*A1783V/WT*^ mice (P28-P42) to evaluate the temperature at which the mice seized. A rectal probe was inserted 10 min before the time-to-peak effect (TPE) of the test drug(s) and hyperthermia-induced seizure test. Before the test started, mice acclimated to the temperature probe for 5 min in a glass chamber. Mice were placed under a heat lamp, and the core temperature was gradually elevated by 1°C every 2 min in the chamber until a generalized seizure was observed or the temperature reached 42.5°C.^18-20^ If no seizure occurred, the mouse was considered seizure-free. A neonatal mouse/rat rectal probe (Braintree Scientific, Inc, Braintree, MA) coupled to a TCAT-2LV controller (Physitemp Instruments, Inc) was used to monitor the core body temperature. Male and female *Scn1a*^*A1783V/WT*^ mice were randomly assigned to treatment groups before the test, and observers were blinded to the treatment. A mouse was removed from the study if it had a behavioral seizure between drug administration and testing. The temperature at which mice seized was recorded.

### Materials

CLB and VPA were purchased as reference powders from Sigma-Aldrich. NCLB and VPA-^13^C_6_ were purchased as 1.0 mg/mL stock solutions in MeOH, while CLB-^13^C_6_ and NCLB-^13^C_6_ were purchased as 0.1 mg/mL stock solutions in acetonitrile, all from Sigma Aldrich. CBD was purchased from Cayman Chemical (Ann Arbor, MI). FFA was obtained from Axon Medchem (Groningen, Netherlands). STP and STP-d_9_ were purchased as reference powders from Toronto Research Chemicals (Toronto, Canada)

### Drug dose and administration

All administered drugs were prepared as 0.5% methylcellulose (Sigma) suspensions. All drugs [CLB (5 mg/kg), VPA (75 mg/kg), STP (30 mg/kg, 70 mg/kg, 100 mg/kg, 130 mg/kg), CBD (70 mg/kg, 100 mg/kg, 135 mg/kg, 150 mg/kg)], were administered at 0.01 mL/g volume and tested based on their time of peak effect (TPE) as observed in other models of epilepsy (TPE_CLB, STP, CBD_ = 1 hr, TPE_FFA, LCS_ = 0.5 hr, TPE_VPA_ = 0.25 hr).^17^ Each drug was intraperitoneally (i.p.) administered separately before the hyperthermia-induced seizure test.

### Sample collection

Blood samples were collected from *Scn1a*^*WT/WT*^ mice at various time points: 0.08, 0.17, 0.25, 0.5, 1, 2, 4, and 8-hr post-dose. Mice used in these studies were not subjected to hyperthermia studies. Blood was collected via decapitation into tubes with K_2_EDTA (Becton, Dickson and Company, NJ) as an anticoagulant. Blood was centrifuged (3500xg for 10 min at 4°C) to isolate plasma. Brains were rapidly removed following decapitation and weighed. Whole brains were homogenized in ultrapure water using a probe sonicator to obtain a target brain homogenate concentration of 200 mg/mL. Plasma and brain homogenate samples were stored at −80°C before analysis.

### Bioanalysis

#### CLB, N-CLB, and VPA

CLB was diluted to 1.0 mg/mL in 50:50 methanol:water (MeOH:H_2_O), while VPA was diluted to 10 mg/mL in 50:50 MeOH:H_2_O. NCLB and VPA-^13^C_6_ were purchased as 1.0 mg/mL stock solutions in MeOH, while CLB-^13^C_6_ and NCLB-^13^C_6_ were purchased as 0.1 mg/mL stock solutions in acetonitrile, all from Sigma Aldrich. All mouse plasma samples were diluted 20-fold before extraction, with human plasma with EDTA anticoagulant used as a matrix for calibrators and quality control (QC) samples. Mouse brain homogenate was analyzed without dilution, using mouse brain homogenate as the control matrix for calibrators and QC samples.

A 100 μL aliquot of the 20-fold diluted plasma or undiluted brain homogenate was assayed using a protein precipitation approach. A 50 μL volume of internal standard (400 pg/μL VPA-^13^C_6_, 40 pg/μL CLB-^13^C_6,_ and NCLB-^13^C_6_ in 10:90 MeOH:H_2_O) was added to each sample aliquot, followed by 1 mL of cold acetonitrile. The samples were vortex-mixed and centrifuged to separate the supernatant from the protein pellet. The supernatant was then transferred to a fresh microcentrifuge tube and centrifuged again to ensure the removal of all solid particulates from the sample. The supernatant was transferred into a polypropylene tube and dried under house air (15 psi) in a TurboVap set (40°C). Samples were then reconstituted with 200 μL of 10:90 MeOH:H_2_O, centrifuged, vortex mixed, and transferred to autosampler vials for analysis.

A 10 μL sample volume was injected for analysis on a Waters Acquity UPLC with a Quattro Premier XE UHPLC-MS/MS system. Chromatography was performed using a Phenomenex (Torrance, CA) Luna Omega Polar C18, 1.6 μm (2.1 x 50 mm), and gradient elution was maintained at a 300 μL/min flow rate. Mobile phases consisted of (A) 10mM ammonium acetate (pH 5.5) and (B) methanol. Analytes were monitored using the following mass transitions (collision energy, CE): 287.1⟶245.1 (NCLB, CE=20V); 293.1⟶251.1 (NCLB-^13^C_6_, CE=20V); 301.1⟶259.1 (CLB, CE=20V); 307.1⟶265.1 (CLB-^13^C_6_, CE=20V); 143.1⟶143.1 (VPA, CE=5V); and 149.1⟶149.1 (VPA-^13^C_6_, CE=5V). CLB, NCLB, and their internal standards (IS) utilized positive electrospray ionization (ESI), whereas VPA and its internal standard (IS) used negative ESI. The calibration curves utilized 1/x^2^ weighted linear regression between 4-200 ng/mL for CLB and NCLB and 0.2-10 μg/mL for VPA.

#### STP

STP was diluted to 1.0 mg/mL in 90:10 MeOH:H_2_O. Mouse plasma samples were diluted 100-fold into 50:50 MeOH:H_2_O before analysis. A 200 μL aliquot of the 100-fold diluted sample was analyzed with calibrators, and QC was prepared in 1% mouse plasma in 50:50 MeOH:H_2_O. Mouse brain homogenate was diluted 10-fold into 50:50 MeOH:H_2_O before analysis, and a 200 μL aliquot was assayed against calibrators and QCs prepared in 10% mouse brain homogenate in 50:50 MeOH:H_2_O.

STP was extracted using a basified liquid-liquid extraction procedure in polypropylene tubes. A 50 μL volume of 40 pg/μL STP-d_9_ in 50:50 MeOH:H_2_O was added to each sample. The sample pH was then adjusted with 0.5 mL of 5% (v:v) ammonium hydroxide in type-1 water and extracted with 2.0 mL of methyl tert-butyl ether (MTBE). The sample was mixed well and centrifuged to separate the organic and aqueous layers. The sample was frozen at -80°C for 30 min, and the organic layer was decanted into a fresh tube. The organic layer was dried under house air (15 psi) in a TurboVap set (40°C). The sample was then reconstituted with 200 μL of 50:50 MeOH:H_2_O and transferred to an autosampler vial for analysis.

A 20 μL sample volume was injected onto a Waters Acquity UPLC and autosampler coupled with a ThermoScientific TSQ Quantum Access MS/MS. Chromatography utilized a Phenomenex Luna Omega PS C18 3μm (2.1 x 100mm) column, and gradient elution was maintained at a 300 μL/min flow rate. Mobile phases comprised (A) 0.1% formic acid in type-1 water and (B) methanol. STP and its IS were ionized using positive ESI, and the mass transitions (CE) 217.2⟶187.2 (11V) and 226.2⟶196.2 (11V) were monitored for STP and STP-d_9_, respectively. The assay’s dynamic range was 1-1000 ng/mL, fitting with a 1/x weighted linear regression.

#### CBD and 7-OH-CBD

CBD and 7-OH-CBD were purchased as 1.0 mg/mL stock solutions in MeOH from Sigma-Aldrich, while CBD-d_3_ and 7-OH-CBD-d_3_ were purchased as 0.1 mg/mL stock solutions in MeOH. Plasma and brain homogenate were diluted before analysis 10-fold into human plasma with EDTA anticoagulant. Calibrators and QC samples were also prepared in human plasma. A 200 μL aliquot of the 10-fold diluted sample was analyzed following a method previously validated by the Center for Human Toxicology.

Samples were extracted using a liquid-liquid extraction under acidic conditions in polypropylene tubes. After addition of 50 μL of 20 pg/μL CBD-d_3_ and 7-OH-CBD-d_3_ in 50:50 MeOH: H_2_O to each sample, the sample was acidified using 0.5 mL of 2 mM ammonium formate (pH 3.5) and extracted using 2.0 mL of 5:1 hexane:MTBE. The sample was mixed and centrifuged. The sample was frozen at -80°C for 30 min, and the organic layer was decanted into a fresh tube. The organic layer was dried under house air (15 psi) in a TurboVap set (40°C). The sample was then reconstituted with 200 μL of 50:50 MeOH:H_2_O and transferred to an autosampler vial for analysis.

A 30 μL sample volume was injected onto a ThermoScientific Accela LC pump, and an autosampler interfaced with a ThermoScientific TSQ Vantage MS/MS. Chromatography used a Phenomenex Luna Omega PS C18, 3 μm (2.1 x 100 mm) column, and gradient elution maintained at a 250 μL/min flow rate. Mobile phases consisted of (A) 1 mM ammonium formate (pH 3.5) and (B) methanol. Positive ESI was used for CBD, 7OH-CBD, and their IS, monitoring the mass transitions (CE): 315.2⟶193.2 (20V, CBD), 318.2⟶196.2 (20V, CBD-d_3_), 331.2⟶313.2 (10V, 7OH-CBD), and 334.2⟶316.2 (10V, 7OH-CBD-d_3_). Calibration curves for both analytes are linearly regressed with 1/x^2^ weighting between the concentrations of 0.5-100 ng/mL.

### PK Analysis

Before PK analyses, concentrations were averaged at each time point for each combination of drug regimen, tissue, and sample time. PK analyses were conducted in Phoenix WinNonLin v8.3, using the linear trapezoidal linear interpolation option calculation method in the non-compartmental analysis module. The software was allowed to determine the best-fit line for the elimination slope for all calculations. The parameters of interest for this analysis were the area under the concentration-time curve from zero to infinity (AUC_0-inf_), maximum concentration (C_max_), and half-life (t_1/2_) for CLB, N-CLB, STP, CBD, and 7-OH-CBD.

### Statistical analysis

Statistical analyses were conducted using a Log-rank (Mantel-Cox) test for the hyperthermia-induced test. A power analysis used α=0.05 and the hazard ratio=0.20. Significance was defined as a p-value<0.05. All analysis was conducted with GraphPad Prism 9.0. Data are presented as mean±SD.

## Results

### Higher doses of STP in combination with CLB+VPA were effective against hyperthermia-induced seizures in *Scn1a*^*A1783V/WT*^ mice

Previously published experiments in our lab determined that STP (100 mg/kg) monotherapy and CLB (5 mg/kg)+VPA (75 mg/kg) were ineffective against hyperthermia-induced seizures in the *Scn1a*^*A1783V/WT*^ mice.^17^ Therefore, in the present experiments, we determined if STP was effective using a clinically relevant combination with both CLB and VPA. Different doses of STP were co-administered with CLB and VPA, and the seizing temperatures were determined and compared to the temperature at which seizures occurred in a control group receiving CLB (5 mg/kg) and VPA (75 mg/kg) combination without STP. STP [(100 mg/kg; median seizing temp -40.4°C ^**^p=0.0471) or (130 mg/kg; median seizing temp -40.2°C ^*^p=0.0039)] + CLB (5 mg/kg) + VPA (75 mg/kg) treated mice had significantly higher temperature thresholds than CLB (5 mg/kg) + VPA (75 mg/kg) treated *Scn1a*^*A1783V/WT*^ mice (median seizing temps -39.35°C and 39.15 °C respectively). **(Figure 1)**. Lower doses of STP (30 and 70 mg/kg) added to CLB+VPA did not significantly change temperature thresholds compared to CLB+VPA without STP. Additionally, signs of sedation were observed in CLB+VPA+STP (130 mg/kg)-treated mice but not in the CLB+VPA-treated group. Data from the CLB+VPA+STP (30 mg/kg and 100 mg/kg) hyperthermia-induced seizure test was reprinted with permission from Pernici et al.^17^

**Figure 1:**
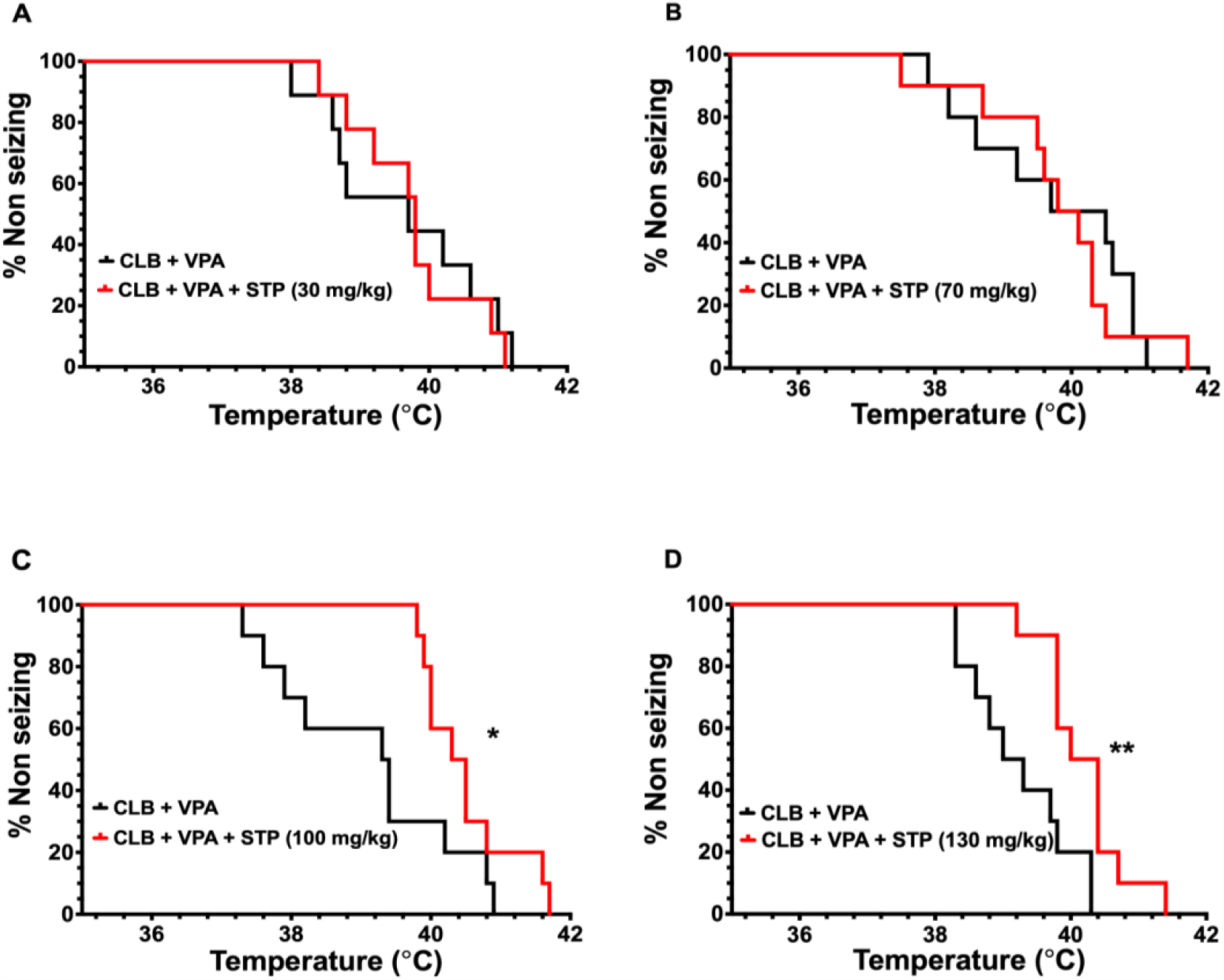
Seizure protection profile of add-on therapy of STP to CLB and VPA in *Scn1a*^*A1783V/WT*^ mice. (A) STP (30 mg/kg) + CLB (5 mg/kg) + VPA (75 mg/kg) did not significantly affect the temperature at which mice seized (CLB + VPA + STP: n = 9, CLB + VPA: n = 9 p = .7973) (B) STP (70 mg/kg) + CLB (5 mg/kg) + VPA (75 mg/kg) failed to over any significant protection against hyperthermia-induced seizures in mice (CLB + VPA + STP: n = 10, CLB + VPA: n = 10 p = .7743) (C) Add-on STP (100 mg/kg) to CLB (5 mg/kg) + VPA (75 mg/kg) significantly increased the median seizing temperature threshold at which mice seize (CLB + VPA + STP: n = 10, CLB + VPA: n = 10) (D) Add-on STP (130 mg/kg) to CLB (5 mg/kg) + VPA (75 mg/kg) significantly increased the median seizing temperature threshold at which mice seize (CLB + VPA + STP: n = 10, CLB + VPA: n = 10). *p < .05, ^**^p < 0.005; Log-rank (Mantel-Cox). Figures 1A and 1C were modified and published with permission from *(Pernici CD et al*., *2021)*.

### Plasma and brain PK profiles demonstrate potential drug-drug interactions between STP, CLB, and VPA

Treatment with CLB (5mg/kg)+VPA (75mg/kg)+STP (100mg/kg) significantly increased the temperature threshold for seizure induction in *Scn1a*^*A1783V/WT*^ mice, indicating a protective effect of this drug combination. Thus, brain and plasma PK evaluation of CLB, N-CLB, VPA, and STP was performed in wild-type mice from our breeding colony to determine brain and plasma exposure sufficient for drug target engagement, inform changes in analyte concentrations due to DDI, and guide optimal dosing strategies for sub-chronic studies. Brain PK profiles were generated for CLB, N-CLB, VPA, and STP in stand-alone or combination therapies (**Figure 2** and **Table 1)**. STP exposure in the triple-drug therapy was higher than STP monotherapy [AUC_0-inf,b_15.1 μg^*^hr/g vs AUC_0-inf,b_ 5.56 μg*hr/g]. STP C_max_ in plasma and brain in STP+CLB+VPA treatment were C_max,p_ =30.7 μg/mL = C_max,b_ =5.34 μg/g, respectively. CLB exposures in the brain were increased in animals co-treated with STP [AUC_0-inf,b_ 7.69 μg*hr/g] compared to CLB-only treatment [AUC_0-inf,b_ 0.744 μg^*^hr/g] or CLB+VPA co-administration [AUC_0-inf,b_ 0.637 μg^*^hr/g]. Total N-CLB exposures in the brain were higher in triple-drug-treated mice than in CLB+VPA or CLB monotherapy. [AUC_0-inf,b_ 16.3 μg^*^hr/g vs AUC_0-inf,b_ 11.6 μg^*^hr/g or AUC_0-inf,b_ 5.13 μg^*^hr/g]. **(Table 1**).

**Table 1:** PK parameters following intraperitoneal administration of CLB, VPA, STP, and CBD in plasma and brain of adult *Scn1a*^*WT/WT*^ DS mice.

| Treatment type | Analytes | Plasma |  |  | Brain |  |  |
| --- | --- | --- | --- | --- | --- | --- | --- |
|  |  | PK Parameters |  |  |  |  |  |
|  |  | AUC (µg*hr/mL) | Cmax (µg/mL) | t1/2 (hr) | AUC (µg*hr/g) | Cmax (µg/g) | t1/2 (hr) |
| Single Drug | CLB | 0.648 | 0.418 | 1.86 | 0.749 | 0.446 | 2.46 |
|  | N-CLB | 5.53 | 0.77 | 5.29 | 11.6 | 0.966 | 10.4 |
|  | VPA | 129 | 155 | 0.397 | 12.3 | 23.6 | 0.28 |
|  | STP | 53.3 | 34.3 | 0.868 | 5.56 | 4.39 | 0.863 |
|  | CBD | 4.21 | 1.39 | 2.25 | 5.43 | 1.42 | 2.74 |
|  | 7-OH-CBD | 3.3 | 0.67 | 3.32 | 2.71 | 0.62 | 3.25 |
| Double (CLB + VPA) | CLB | 0.462 | 0.54 | 1.530 | 0.637 | 0.76 | 0.904 |
|  | N-CLB | 3.61 | 0.567 | 2.610 | 5.13 | 0.788 | 3.300 |
|  | VPA | 224 | 243 | 0.623 | 25.1 | 42.5 | 0.469 |
| Triple (CLB + VPA + STP) | CLB | 5.64 | 1.56 | 1.51 | 7.69 | 2.12 | 1.34 |
|  | N-CLB | 7.13 | 0.88 | 3.83 | 16.3 | 1.22 | 11.2 |
|  | VPA | 209 | 166 | 0.67 | 27.1 | 35.5 | 0.274 |
|  | STP | 103 | 30.7 | 1.51 | 15.1 | 5.34 | 1.24 |
| Triple (CLB + VPA + CBD) | CLB | 2.05 | 0.579 | 2.11 | 2.87 | 0.858 | 1.69 |
|  | N-CLB | 14.8 | 0.76 | 13 | 23.2 | 0.91 | 17.1 |
|  | VPA | 295 | 233 | 1.55 | 36.1 | 42.1 | 1.19 |
|  | CBD | 6.29 | 1.24 | 2.78 | 12.9 | 1.33 | 5.84 |
|  | 7-OH-CBD | 5.86 | 0.72 | 4.28 | 5.97 | 0.622 | 5.99 |

Abbreviations: AUC, area under the concentration–time curve; CLB, clobazam; Cmax, maximum concentration; CBD, cannabidiol; n/a, not available; PK, pharmacokinetic; STP, stiripentol; t1/2, half-life; VPA, sodium valproate.

**Figure 2:**
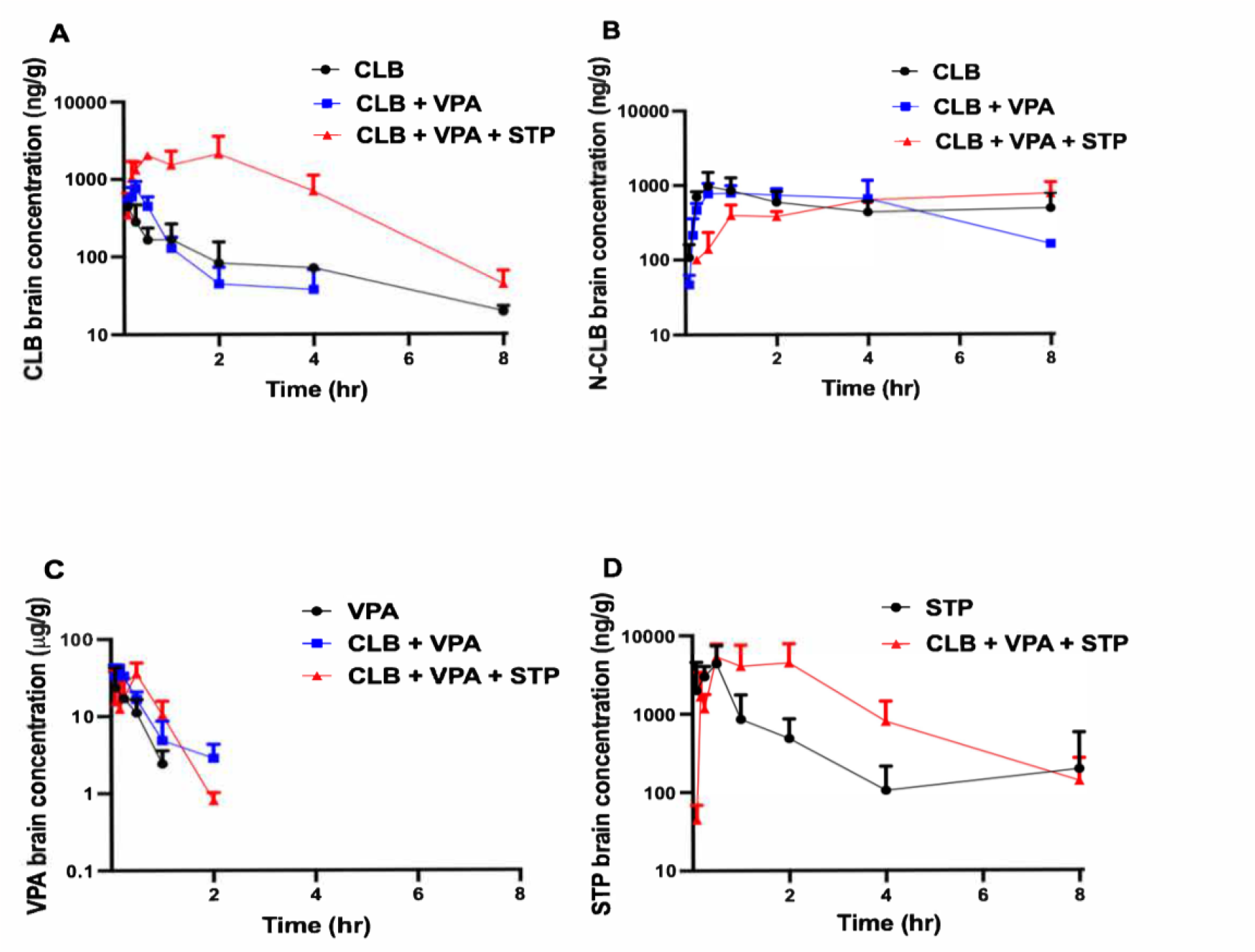
Concentration–time curves of (A) CLB (5 mg/kg), (B) N-CLB, (C) VPA (75 mg/kg), and (D) and STP (100 mg/kg) in the brain of adult *Scn1a*^*WT/WT*^ mice following intraperitoneal administration. Brain samples were obtained 0– 8 hr post-dose. Each data point is the mean brain concentration value for four mice.

### Higher doses of CBD add-on to CLB and VPA were effective against hyperthermia-induced seizures in *Scn1a*^*A1783V/WT*^ mice

Previous experiments in our lab demonstrated that when CBD (135 mg/kg) was administered alone, it failed to increase temperature thresholds in *Scn1a*^*A1783V/WT*^ mice.^17^ In this study, we assessed whether different doses of CBD added to CLB+VPA would be effective against hyperthermia-induced seizures in *Scn1a*^*A1783V/WT*^ mice. Various doses of CBD (70 mg/kg, 100 mg/kg, 135 mg/kg, or 150 mg/kg) in combination with CLB (5 mg/kg)+VPA (75 mg/kg) were tested to determine its activity against hyperthermia-induced seizure. CBD [135 mg/kg; median seizing temp -40.7°C ^*^p=0.027) or (150 mg/kg; median seizing temp -39.7°C ^*^p=0.01] + CLB+VPA showed a significant increase in threshold temperature to induce seizures when compared to CLB (5 mg/kg) + VPA (75 mg/kg) treated *Scn1a*^*A1783V/WT*^ mice **(Figure 3)**. Lower doses of CBD (70 and 100 mg/kg) added to CLB+VPA did not change temperature thresholds compared to CLB+VPA without CBD. Signs of hypothermia were observed in all doses of CLB+VPA+CBD-treated mice but not in the CLB+VPA-treated group.

**Figure 3:**
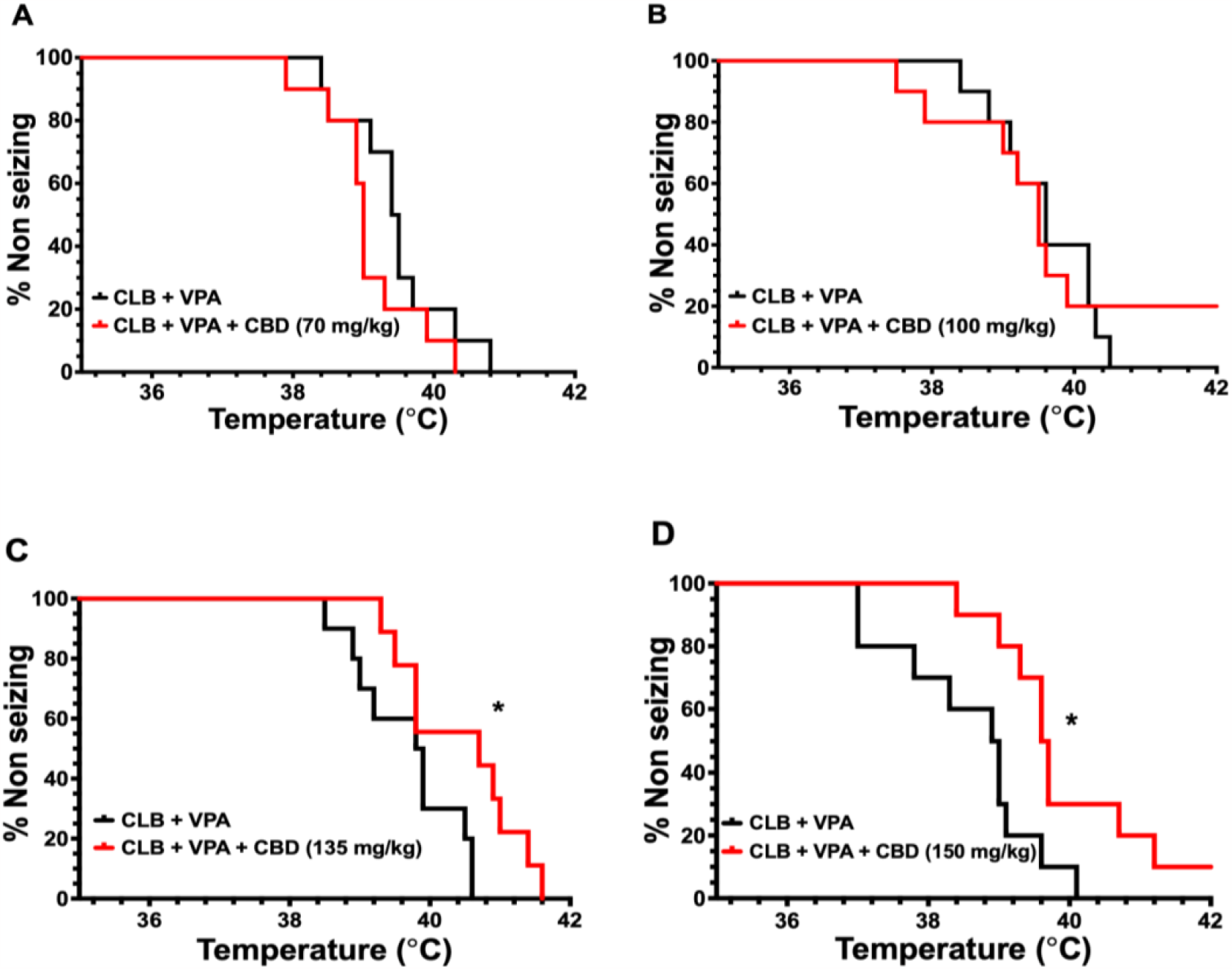
Antiseizure efficacy of CBD add-on to CLB and VPA in *Scn1a*^*A1783V/WT*^ mice. (A) CBD (70 mg/kg) + CLB (5 mg/kg) + VPA (75 mg/kg) did not significantly protect mice against hyperthermia-induced seizures (CLB + VPA + CBD: n = 10, CLB + VPA: n = 10 p = .7973) (B) CBD (100 mg/kg) + CLB (5 mg/kg) + VPA (75 mg/kg) failed to over any significant protection against hyperthermia-induced seizures in mice (CLB + VPA + CBD: n = 10, CLB + VPA: n = 10 p = .8746) (C) Add-on CBD (135 mg/kg) to CLB (5 mg/kg) + VPA (75 mg/kg) offered protection against hyperthermia-induced seizures in mice seize (CLB + VPA + CBD: n = 9, CLB + VPA: n = 10) (D) Add-on CBD (150 mg/kg) to CLB (5 mg/kg) + VPA (75 mg/kg) offered protection against hyperthermia-induced seizures in mice seize (CLB + VPA + CBD: n = 10, CLB + VPA: n = 10) ^*^p < .05; Log-rank (Mantel-Cox).

### Plasma and brain PK profiles demonstrate potential DDIs between CBD, CLB, and VPA

Treatment with CLB(5mg/kg)+VPA(75mg/kg)+CBD(135mg/kg) was effective against hyperthermia-induced seizures in *Scn1a*^*A1783V/WT*^ mice. Therefore, brain and plasma concentrations of CBD, CLB, N-CLB, and VPA were analyzed to determine whether the effective doses resulted in brain and plasma concentrations within the human therapeutic range to ensure an accurate assessment of relevant drug responses in our DS model. The PK parameters of CBD treatment with CLB, N-CLB, and VPA in the brains of mice are displayed **in Figure 4 and Table 1**. CBD brain exposure was higher in the triple-drug treatment group when compared to CBD monotherapy [AUC_0-inf,b_ 12.8 μg*hr/g vs AUC_0-inf,b_ 5.43 μg^*^hr/g]. CBD administration increased the exposure of CLB in the brain when compared to CLB in CLB-alone or CLB+VPA co-treatments [AUC_0-inf,b_ 2.87 μg^*^hr/g vs AUC_0-inf,b_ 0.744 μg^*^hr/g or AUC_0-inf,b_ 0.637 μg^*^hr/g]. CBD increased N-CLB exposure when combined with CLB and VPA compared to the N-CLB concentrations in CLB-alone or CLB+VPA treatments [AUC_0-inf,b_ 23.2 μg^*^hr/g vs AUC_0-inf,b_ 1.16 μg^*^hr/g or AUC_0-inf,b_ 5.13 μg*hr/g]. CBD increased the exposure of brain VPA in CBD+CLB+VPA compared to CLB-alone or CLB+VPA treatments [AUC_0-inf,b_ 36.1 μg*hr/g vs AUC_0-inf,b_ 12.3 μg*hr/g or AUC_0-inf,b_ 25.1 μg*hr/mL] respectively.

**Figure 4:**
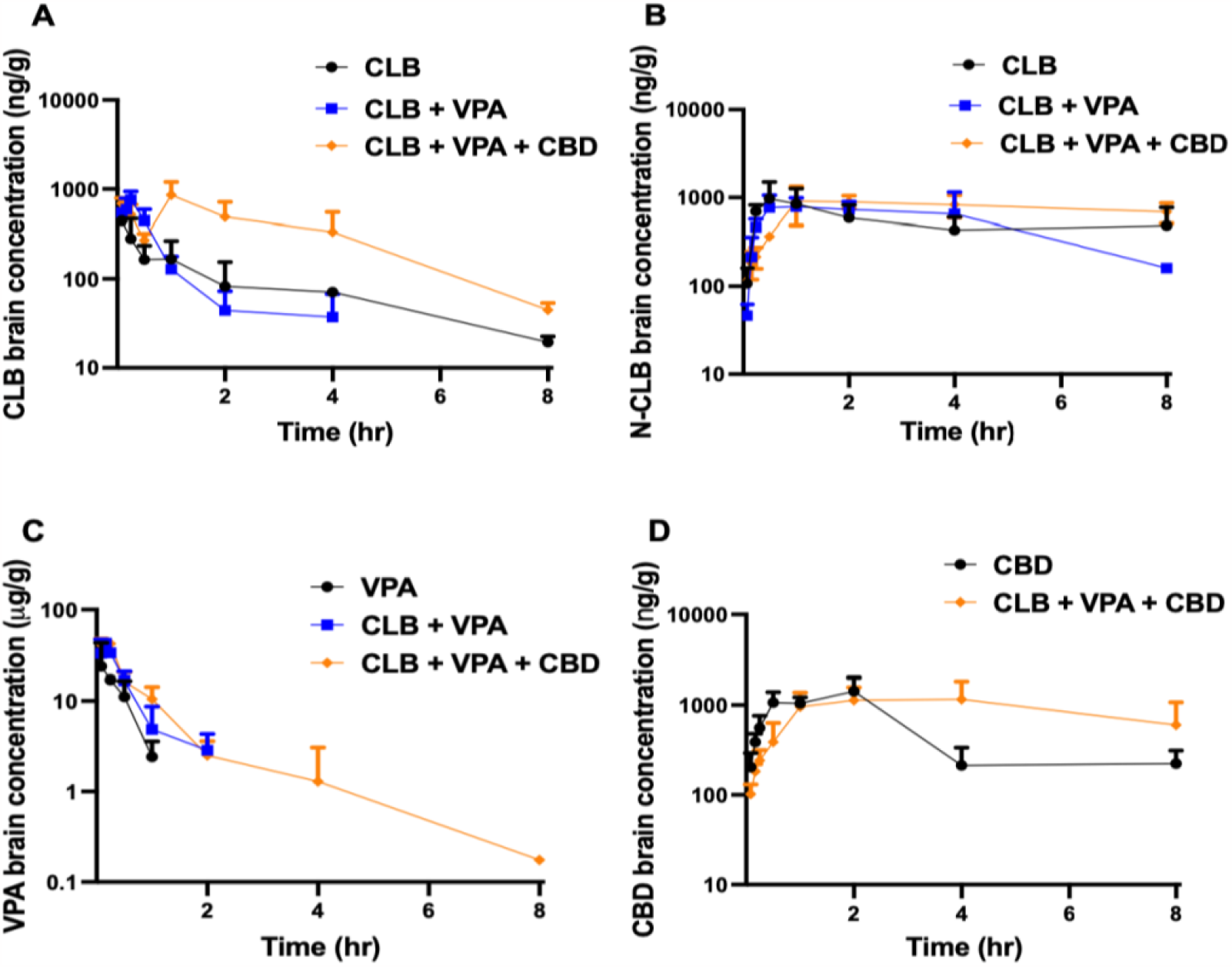
Concentration–time curves of (A) CLB (5 mg/kg), (B) N-CLB, (C) VPA (75 mg/kg), and (D) and CBD (135 mg/kg) in the brain of adult *Scn1a*^*WT/WT*^ mice following intraperitoneal administration. Brain samples were obtained 0– 8 hr post-dose. Each data point is the mean brain concentration value for four mice.

### FFA and LCS treatment as adjunctive therapies to CLB and VPA were not effective against hyperthermia-induced seizures in *Scn1a*^*A1783V/WT*^ mice

Similar to STP and CBD, as previously published, other FDA-approved and investigational compounds for treating DS, such as FFA and LCS, were insufficient to prevent hyperthermia-induced seizures in the Scn1aA1783V/WT mice when delivered alone.^17^ Therefore, we evaluated the effect of FFA or LCS administration in combination with CLB+VPA against hyperthermia-induced seizures in DS mice. At all doses of FFA (5, 10, 20, and 30 mg/kg) administered in combination with CLB (5 mg/kg) + VPA (75 mg/kg), the combination treatment failed to confer protection from hyperthermia-induced seizures in the *Scn1a*^*A1783V/WT*^ mice. Similarly, there were no significant differences in the seizing temperatures when multiple doses of LCS (5, 7, 10, and 15 mg/kg) were combined with CLB (5 mg/kg) + VPA (75 mg/kg) compared to the control-treated group of CLB (5 mg/kg) + VPA (75 mg/kg) (p>0.05) **(Table 2)**. Since CLB+VPA+FFA and CLB+VPA+LCS were ineffective against hyperthermia-induced seizures, we did not follow up with PK studies to determine DDI between the compounds.

**Table 2:** Summary of the efficacy of STP, CBD, FFA, and LCS as add-on therapies to CLB and VPA against hyperthermia-induced seizures following intraperitoneal administration in *Scn1*^*A1783A/WT*^ mice.

| STP - add on |  |  |  |  |
| --- | --- | --- | --- | --- |
| Drug treatment | Seizing temp (°C) | Control treatment | Seizing temp (°C) | Statistical significance (Log-Rank) |
| CLB (5 mg/kg) + VPA (75 mg/kg) + STP (30 mg/kg) | 39.8 (n =9) | CLB (5 mg/kg) + VPA (75 mg/kg) | 39.7 (n= 9) | p = 0.797 |
| CLB (5 mg/kg) + VPA (75 mg/kg) + STP (70 mg/kg) | 39.5 (n =10) |  | 39.7 (n= 9) | p = 0.797 |
| CLB (5 mg/kg) + VPA (75 mg/kg) + STP (100 mg/kg) | 40.4 (n =10) |  | 39.35 (n= 10) | *p = 0.047 |
| CLB (5 mg/kg) + VPA (75 mg/kg) + STP (130 mg/kg) | 40.2 (n =10) |  | 39.15. (n= 10) | **p = 0.004 |
| CBD - add on |  |  |  |  |
| Drug treatment | Seizing temp (°C) | Control treatment | Seizing temp (°C) | Statistical significance (Log-Rank) |
| CLB (5 mg/kg) + VPA (75 mg/kg) + CBD (70 mg/kg) | 39.0 (n =10) | CLB (5 mg/kg) + VPA (75 mg/kg) | 39.5 (n= 10) | p = 0.186 |
| CLB (5 mg/kg) + VPA (75 mg/kg) + CBD (100 mg/kg) | 39.5 (n =10) |  | 39.6 (n= 9) | p = 0.875 |
| CLB (5 mg/kg) + VPA (75 mg/kg) + CBD (135 mg/kg) | 40.7 (n =9) |  | 39.85 (n= 10) | *p = 0.027 |
| CLB (5 mg/kg) + VPA (75 mg/kg) + CBD (150 mg/kg) | 39.7 (n =10) |  | 38.95 (n= 10) | *p = 0.01 |
| FFA - add on |  |  |  |  |
| Drug treatment | Seizing temp (°C) | Control treatment | Seizing temp (°C) | Statistical significance (Log-Rank) |
| CLB (5 mg/kg) + VPA (75 mg/kg) + FFA (5 mg/kg) | 39.4 (n =10) | CLB (5 mg/kg) + VPA (75 mg/kg) | 39.4 (n= 10) | p = 0.787 |
| CLB (5 mg/kg) + VPA (75 mg/kg) + FFA (10 mg/kg) | 39.4 (n =10) |  | 39.0 (n= 10) | p = 0.334 |
| CLB (5 mg/kg) + VPA (75 mg/kg) + FFA (20 mg/kg) | 39.0 (n =10) |  | 38.4 (n= 10) | p = 0.982 |
| CLB (5 mg/kg) + VPA (75 mg/kg) + FFA (30 mg/kg) | 39.2 (n =10) |  | 39.1 (n= 10) | p = 0.345 |
| LCS - add on |  |  |  |  |
| Drug treatment | Seizing temp (°C) | Control treatment | Seizing temp (°C) | Statistical significance (Log-Rank) |
| CLB (5 mg/kg) + VPA (75 mg/kg) + LCS (5 mg/kg) | 39.6 (n =10) | CLB (5 mg/kg) + VPA (75 mg/kg) | 39.9 (n= 10) | p = 0.346 |
| CLB (5 mg/kg) + VPA (75 mg/kg) + LCS (7 mg/kg) | 39.5 (n =10) |  | 39.7 (n= 9) | p = 0.513 |
| CLB (5 mg/kg) + VPA (75 mg/kg) + LCS (10 mg/kg) | 38.3 (n =10) |  | 39.7 (n= 9) | p = 0.939 |
| CLB (5 mg/kg) + VPA (75 mg/kg) + LCS (15 mg/kg) | 38.9 (n =10) |  | 39.4 (n= 10) | p = 0.133 |

Abbreviations: CBD, cannabidiol, CLB, clobazam; FFA, fenfluramine; LCS, lorcaserin; STP, stiripentol, VPA, sodium valproate.

## Discussion

Although newer drugs have been approved for treating patients with DS, pharmacoresistance remains common in DS.^21^ Thus, there is a critical need to introduce novel drugs and better treatment strategies for DS. DS is usually treated with multiple ASMs since monotherapy is generally inadequate.^22^ Using etiologically relevant and well-characterized preclinical models that include comprehensive PK profiling incorporation is helpful for clinical studies. Furthermore, comparison to drug exposures in the human therapeutic range is essential for translating ASMs into preclinical success.^23^ The present study used single administration (i.p) doses and a polytherapy paradigm that demonstrated efficacy against hyperthermia-induced seizures. We showed alterations in some PK parameters such as AUC, C_max,_ and half-life, suggesting potential DDI between STP, CBD, CLB, and VPA that could underlie the antiseizure effects observed in the triple-drug treatment studies. This sets the stage to guide the design of a dosing regimen for a triple-drug treatment in a mouse model of DS for sub-chronic spontaneous seizure drug screening.

Treatment with STP (100 mg/kg and 130 mg/kg) combined with both CLB+VPA demonstrated efficacy against hyperthermia-induced seizures in *Scn1a*^*A1783V/WT*^ mice compared to the CLB+VPA treatment group. The PK profiles of STP showed that the peak plasma concentration of the effective dose of STP as an adjunctive therapy was 30.7 μg/mL, above the human therapeutic plasma concentration range of 8-16 μg/mL, similar to reports by Jullien et al.^24^ In contrast, Cao et al.^25^ reported that STP was effective as monotherapy at high doses (150-350 mg/kg) in a hyperthermia-induced seizure test in the *Scn1a*^*RX/+*^ mice, suggesting that doses higher than 100 mg/kg in STP stand-alone treatment may be required to demonstrate efficacy against hyperthermia-induced seizures in a DS mouse model. Other experiments also suggest that the antiseizure effects of STP are potentiated by the interaction with other ASMs, such as CLB, via DDIs.^26^ We observed greater total brain STP exposure when STP is combined with CLB and VPA compared to STP alone treatment. This suggests that co-administering STP with CLB and VPA might enhance the absorption of STP compared to STP monotherapy. Furthermore, a trend of more sustained peak brain concentrations of STP in the triple-drug group was also observed compared to the STP monotherapy. This phenomenon could partly be attributed to a perfusion-limiting process at the blood-brain barrier, potentially due to CLB and VPA. CLB has been reported to increase STP concentration by 37%; however, the potential mechanism responsible for the increase of STP levels by CLB remains unclear.^27^ Additionally, VPA inhibits CYP 3A4 activity, an enzyme that mediates the first-pass effect of STP, increasing STP exposure.^28^ STP elimination kinetics remains controversial, and data about the influence of other ASMs, such as CLB and VPA, on STP is sparse^27,29^ In the present study, although we observe a trend of a slightly longer half-life of STP in the triple-drug group compared to the STP stand-alone, the short duration (8-hr maximum time-point) of our study made it challenging to make strong inferences from the observed STP half-lives. Nevertheless, our study’s elimination phase of STP displayed a multiphasic profile similar to that reported by Levi et al.^30^ Overall, we found that plasma and brain STP exposures in mice increase when co-administered with CLB and VPA, consistent with human studies.

When CBD (135 mg/kg and 150 mg/kg) was combined with CLB and VPA treatment, a significant shift in the temperature at which hyperthermia-induced seizures in S*cn1a*^*A1783V/WT*^ mice was observed compared to CLB plus VPA treatment. Several studies have reported that CBD (100-200 mg/kg) demonstrates an antiseizure effect when given alone or co-administered with CLB^20,31^ Potential DDI, coupled with differences in the *Scn1a* gene mutation types and independent PD effects of CLB and CBD, may contribute to the differences in observed outcomes when CBD is given alone or combined with CLB.^20,32^ We observed a two-fold increase in brain exposure to CBD in the triple-drug treatment group of CBD+CLB+VPA compared to CBD monotherapy. The higher AUC values observed for CBD in combination therapy could be associated with DDI with CLB and/or VPA that reduces CBD clearance. CYP 2C19 and CYP 3A4 catalyze the metabolism of CBD.^33^ VPA is a mild CYP 2C19 and CYP 3A4 inhibitor.^34^ Additionally, CBD is a known substrate of CYP 2D6, which CLB extensively inhibits.^35,36^ This coordinated inhibition of CYP450 enzymes relevant to CBD metabolism by CLB and VPA is expected to slow CBD clearance, increasing CBD half-life and total exposure under those conditions. Moreover, CBD is a significant substrate of drug efflux transporter protein, breast cancer resistance protein (BCRP), located at the blood-brain barrier.^37^ VPA inhibits the transport of BCRP substrates such as CBD, thus reducing the efflux of CBD from the brain and increasing CBD brain concentrations.^38^ Our study demonstrated changes in CBD brain PK profiles when CBD is added to CLB and VPA, thus suggesting an alteration in the dosage regimen of CBD when given concurrently with CLB and VPA for sub -chronic studies.

There were slight differences in the PK parameters of CLB and N-CLB in mice treated with CLB alone compared to CLB in combination with other drugs, suggesting potential DDI between CLB and VPA, STP, and CBD.^20,39^ CLB N-demethylation to N-CLB is the main metabolic pathway for CLB^40,41^ Higher CLB exposure and C_max_ values were observed when CLB was co-administered with VPA and STP or CBD compared to CLB+VPA treatment. This could be attributed to the increased absorption of CLB into the plasma and brain. CLB is a substrate of the P-gp^36^, while STP and CBD are P-gp inhibitors and, thus, may reduce the efflux of CLB from the brain.^42-44^ Our data indicate higher N-CLB (> 2-fold increase) concentrations and slightly longer half-lives were observed in triple-drug therapies compared to CLB+VPA treatment. Additionally, higher CLB and N-CLB exposures and longer half-lives were observed in CBD add-on compared to STP add-on, suggesting differences in the extent of interactions between STP or CBD and CLB and N-CLB. CLB is known to increase VPA exposure slightly.^45^ This study demonstrates significant DDI between CLB and N-CLB with STP or CBD that could contribute to the overall observed efficacy in the triple-drug treatment groups.

The present study also evaluated the efficacy of FFA for treating seizures associated with DS as an adjunctive drug. At all doses tested, FFA failed to protect against hyperthermia-induced seizures when added to CLB and VPA. Although several preclinical and clinical studies have reported the successful use of FFA as an add-on treatment for DS, not all patients with DS have their seizures adequately suppressed by FFA, and not all FFA-treated DS mice are protected from hyperthermia-induced seizures.^46-48^ Like FFA, LCS failed to protect against seizures when added to CLB and VPA in *Scn1a*^*A1783V/WT*^ mice. Considering this study was an acute hyperthermia-induced seizure test with compounds administered simultaneously, a PK study of these compounds might help determine appropriate doses sufficient to provide sufficient therapeutic plasma and brain levels. Nevertheless, this suggests that our model is a drug-resistant DS model for screening novel compounds for DS.

There are limitations of the present study. Each time point of the PK studies is pooled data from four WT mice rather than longitudinal samples collected from the same animal over time. This approach impairs the determination of inter-animal variability in PK, preventing hypothesis testing of PK differences observed in the study. However, this approach is necessary to permit brain sampling that provides critical data on drug PK at the site of action. Moreover, the additional volume in the intraperitoneal space may alter perfusion in the triple-drug studies compared to a single drug. Sampling through 8 hrs also limited robust determination of half-life for some compounds. Additionally, we did not perfuse brain tissue with saline before homogenization; thus, residual blood in the sample may influence the measured brain concentrations. Nevertheless, the work reported here demonstrates the importance of determining DDI using polypharmacy approaches.

To find effective therapies for DS, investigational compounds may also need to be screened against spontaneous seizures. Thus, PK data from this study will guide the design of appropriate sub-chronic dosage regimens for evaluating the efficacy of STP or CBD when co-administered with CLB and VPA against spontaneous seizures. The ETSP contract site utilizes the hyperthermia-induced seizure test in the DS model and is essential for identifying other investigational drugs that may be effective when combined with CLB and VPA, similar to clinical practice.

## Acknowledgments

The authors thank the Epilepsy Therapy Screening Program at the National Institute of Neurological Disorders and Stroke for review and comments on the manuscript.

## Author contributions

Study design and results interpretation: JAM, CAR, JER, CSM, and KSW. Data Collection: JAM, KJ, TF, JER. Data analysis: JAM, JER, CSM, and KSW. Manuscript preparation and editing: JAM, JER, CSM, and KSW.

## Data availability

Data from the study would be made available to other investigators for the purpose of replication and re-use upon request.

